# Permethylation as a Strategy for High Molecular Weight Polysaccharide Structure Analysis by NMR – Case Study of *Xylella fastidiosa* EPS

**DOI:** 10.1101/2023.04.24.538115

**Authors:** Ikenna E. Ndukwe, Ian Black, Claudia A. Castro, Jiri Vlach, Christian Heiss, Caroline Roper, Parastoo Azadi

## Abstract

Current practices for structure analysis of extremely large molecular weight polysaccharides via solution-state NMR spectroscopy incorporate partial depolymerization protocols that enable polysaccharide solubilization in suitable solvents. Non-specific depolymerization techniques utilized for glycosidic bond cleavage, such as chemical degradation or ultrasonication, potentially generate structure fragments that can complicate the complete characterization of polysaccharide structures. Utilization of appropriate enzymes for polysaccharide degradation, on the other hand, requires prior structure information and optimal enzyme activity conditions that are not available to the analyst working with novel or unknown compounds. Herein, we describe the application of a permethylation strategy that allows the complete dissolution of the intact polysaccharides for NMR structure characterization. This approach is utilized for NMR analysis of *Xylella fastidiosa* EPS, which is essential for the virulence the plant pathogen that affects multiple commercial crops and is responsible for multibillion dollar losses each year.

## Introduction

High molecular weight polysaccharides often present significant analytical challenges due to their poor inherent solubility or in some cases insolubility in solvents. When solubilized, high molecular weight polysaccharides usually form highly viscous solutions in which extensive hydrogen bonding networks between polymer strands result in severe signal broadening, extremely poor spectral resolution, and sensitivity losses that limit structure analysis via solution-state nuclear magnetic resonance (NMR) spectroscopy. In severe cases, the signals become so broad that no structure information can be derived from the NMR spectra (*vide infra*). Previously, structure analysis of high molecular weight polysaccharides has incorporated mild depolymerization strategies to generate smaller fragments that are then solubilized and analyzed. In particular, chemical degradation methods^1,2^ including oxidative cleavage,^3,4^ mild acid hydrolysis^5–9^ and acetolysis,^10^ have been utilized by various research groups to partially or completely determine the structures of a variety of high molecular weight polysaccharides. These methods yield fragments that are the subsets of the polysaccharide structures but are not a true representation of the repeat units, making them unreliable for complete structure analysis. Ultra-sonication^11–14^ and enzymolysis^15^ are two alternative methods that have also been utilized to obtain tractable samples for structure elucidation of polysaccharides. While the former is considered a mild approach, the generation of secondary structures may be unavoidable due to the non-specific nature of glycosidic bond cleavage of the methodology. Enzymolysis, on the other hand, offers a bond-specific depolymerization strategy, but its application is limited to well-developed systems, which is not always the case for analysts that largely work with unknown carbohydrate structures. Carbohydrates, in contrast to their DNA, RNA, and protein counterparts, are not defined by sequence-based templates. They are synthesized through the concerted actions of enzymes and encompass a diverse array of linkages and stereochemical isomers, precluding facile deconstruction into established subunits. Notwithstanding, polysaccharide depolymerization can be a good way to obtain partial polysaccharide sequence information^3^ and thus should be viewed with consideration of this limitation. The ideal scenario would be to utilize data acquired with oligosaccharide fragments from these depolymerization approaches, in tandem with those acquired with methods that afford NMR solution-state analysis of the intact polysaccharide. It should be noted that solid-state NMR (ssNMR) spectroscopy provides a useful technique to study the intact polysaccharides and has been utilized for investigating inter-molecular interactions between cell wall components and/or to determine the molecular architecture cell wall matrices.^16–20^ Unfortunately, the inherent poor spectral resolution and sensitivity of ssNMR, and to a lesser extent its large sample requirement, limits its general utilization for the full structure characterization of complex polysaccharides.

Herein, we propose the utilization of derivatization techniques, in particular permethylation, to completely solubilize the intact polysaccharide in solvents suitable for solution-state NMR characterization, such as chloroform-*d*. Permethylation removes the extensive intra/inter-molecular hydrogen bonding network of polysaccharides, by replacing all hydroxyl groups with methoxy groups. In a previous article, a case was made for the utilization of permethylation strategies for NMR analysis of soluble and insoluble plant cell wall polysaccharides.^21^ It was observed that ^13^C chemical shift changes (Δδ_C_) due to *O*-methylation of polysaccharides were comparable to glycosylation shifts. In addition, the well-known sugar configuration classification, that defines β anomeric proton signals as resonating lower than ∼4.7 ppm and α anomeric proton signals as resonating higher than ∼ 4.7 ppm, was not impacted by changes in the observed ^1^H chemical shifts (Δδ_H_) due to permethylation. Interestingly, the unique ^13^C chemical shift regions of the permethylated polysaccharides did not change significantly (*O*-Me region, 50 – 65 ppm, hydroxymethyl groups, 60 – 75 ppm, ring residue carbons, 70 – 90 ppm and anomeric carbons, 95 – 115 ppm), hence making permethylation strategies^22,23^ amenable to existing structure characterization protocols of complex polysaccharides. The proposed methodology is utilized to characterize the extracellular polysaccharide (EPS) structure of an economically important micro-organism, *Xylella fastidiosa*.

*X. fastidiosa* is implicated as the causal agent of disease in several economically important commercial crops in North and South America,^24^ and the Mediterranean region of Europe,^25^ including Pierce’s disease (PD).^26^ Although the EPS has been shown to be required for biofilm formation, plant virulence, and insect transmission,^27^ its exact composition and structure is currently unknown. Genetic studies suggest that *X. fastidiosa’* EPS structure is likely similar to the closely related *X. campestris* EPS, xanthan gum.^28^ Characterizing the polysaccharide structures from *X. fastidiosa* and other economically important micro-organisms is important, as these polysaccharides dictate their interaction with the environment.^29^ Microbial polysaccharides frequently are very large and can be challenging to structurally analyze by solution state NMR methods – thus highlighting the value of the proposed permethylation strategy for structure analysis.

## Materials and Methods

### EPS isolation

*X. fastidiosa* cells were initially propagated on solid PD3 plates, harvested, and adjusted to OD_600 nm_= 0.5 in XFM medium. This inoculum was diluted in 15 mL of XFM-pectin liquid to a final OD_600 nm_=0.05 and grown at 28 °C, 180 rpm for six days. Subsequently, the OD_600 nm_ was recorded, and cultures were centrifuged at 7,000 rpm at 4°C for 20 minutes. 10 mL of the supernatant were collected and placed in a 50 mL conical tube, mixed with 30 mL cold 95% ethanol, and placed at -80 °C for 30 min then centrifuged at 7,000 rpm at 4 °C for 20 minutes and supernatants were discarded. Pellets were washed 2X with 10 mL of cold 70% ethanol. A steel grinding ball (size ⅜ in., BC Precision) and 10 mL water was added to the EPS pellet. The tubes were placed inside a shaker (200-300 rpm) for 1 hour to resuspend the pellet. The EPS was then dialyzed against 2 L of deionized water overnight at 4 °C using dialysis membrane tubing with a molecular weight cutoff of 1 kDa (Spectra/Por®, part number 132636). The EPS preparation was frozen, shipped on dry ice and stored at -80°C until use.

### Protein removal

The removal of contaminating protein in the *X. fastidiosa* EPS preparation was accomplished by enzymatic digestion using proteinase K (20 U proteinase/ml, Sigma) in 50 mM MgCl buffer (pH 7.5) overnight at 50 °C (10 mg of EPS/ml solution). After overnight digestion, 10 U/ml pronase (Roche) solution was added, and digestion was carried out as above for an additional 12 hours. After the second round of digestion was complete, the sample was dialyzed against deionized water using a 50-kDa dialysis membrane (SpectraPor®) at 4 °C for 3 days, with the water being exchanged each day.

### Permethylation

Permethylation was carried out as described previously.^21^ However, only one round of permethylation was performed. Sample clean up consisted of a two-step procedure. First, DCM was used to extract fully methylated polysaccharides, followed by an ethanol precipitation step to recover incompletely methylated polysaccharides as described below. After sample permethylation, 300 μl of water was added to the sample to quench the reaction and cause phase separation of iodomethane. Then, 2 ml DCM was added, and the sample was briefly vortexed and centrifuged (50 × g for 10 seconds) using a low-speed benchtop centrifuge. The bottom organic layer was removed and placed into a clean tube and 6 mL ethanol was then added to precipitate undermethylated EPS. The sample was briefly vortexed and centrifuged at 3000 × *g* for 10 minutes to pellet, DMSO containing supernatant was discarded and replaced with 1 mL water followed by 6 mL of clean ethanol. The sample was again vortexed and centrifuged at 3000 × *g*. After ethanol removal, the earlier recovered DCM layer was then added to the ethanol pelleted EPS. The entire solution was dried using a flow of nitrogen and then lyophilized to ensure complete removal of water. The dried sample was then re-permethylated as above. After the second round of methylation, the sample was again DCM extracted and ethanol precipitated as described. Finally, a third round of permethylation was performed. After the third round of methylation, the sample was fully soluble in DCM, and no ethanol precipitation was necessary. The DCM layer was washed 5 times with water to ensure complete removal of DMSO. And the final DCM layer was transferred to a new tube and dried.

### NMR data acquisition

NMR data were acquired on a Bruker 600 MHz spectrometer equipped with a 5 mm TXI cryo-probe at probe temperature 45 °C, except otherwise stated. Between 5 - 10 mg of the samples were permethylated and dissolved in 550 μL chloroform-*d* prior to NMR acquisition. The following general parameters were used for NMR data acquisition. Specific parameters used are indicated in the captions of the respective spectra.

### Permethylated Xylella fastidiosa EPS

^1^H NMR experiment (600 Hz spectral width, 8,192 t1 points and 4 scans); COSY experiment (5144.0*5144.0 F1*F2 spectral width, 1024*512 t2*t1 points and 4 scans); HSQC experiment (5144.0*10570.8 F1*F2 spectral width, 1024*512 t2*t1 points, 145 Hz ^1^*J*_CH_ delay and 10 scans); HSQC_TOCSY experiment (5208.3*10570.8 F1*F2 spectral width, 1024*512 t2*t1 points, 145 Hz ^1^*J*_CH_ delay, 18 ms mixing time and 24 scans); HMQC_NOESY experiment (5208.3*10570.8 F1*F2 spectral width, 1024*512 t2*t1 points, 145 Hz ^1^*J*_CH_ delay, 60 ms mixing time and 182 scans); HMBC experiment (5319.1*10570.8 F1*F2 spectral width, 1536*512 t2*t1 points at 50 % non-uniform sampling in the indirect dimension, 8 Hz ^1^*J*_CH_ delay and 540 scans); TOCSY experiment (6250.0*6250.0 F1*F2 spectral width, 1536*512 t2*t1 points, 60 ms spin-lock time and 8 scans); ROESY experiment (6250.0*6250.0 F1*F2 spectral width, 1536*512 t2*t1 points, 70 ms mixing time and 24 scans).

### TFA hydrolyzate of intact EPS

Quantitative ^1^H NMR spectrum of TFA-hydrolyzed EPS was acquired at 25 °C using 11900 Hz spectral width, 16384 points, four scans and a total relaxation delay of 63 s between each scan. The reference spectra of glucurono-6,3-lactone and glucose/glucuronic acid mixed standard were acquired with the same parameters.

## Results and Discussion

*X. fastidiosa* EPS presented a unique challenge that necessitated a revision of our traditional protocol for structure analysis of high molecular weight polysaccharides. Although somewhat soluble in water,

*X. fastidiosa* EPS gave very broad ^1^H NMR signals that precluded any meaningful structure analysis – even after two attempts at depolymerization via mild acid hydrolysis (**Figure 1**, bottom). In contrast, ^1^H NMR data acquired with the permethylated sample provided improved spectral resolution that revealed some key details about the structure of the EPS (**Figure 1**, top). This improved spectral resolution is attributed to the elimination of intra- and inter-molecular hydrogen bonding interactions that limited the rotational freedom of the native polysaccharides, as hydroxyl groups are replaced by *O*-methyl groups, in combination to the utilization of a less viscous solvent like chloroform-*d* for NMR data acquisition. Each polysaccharide strand is able to diffuse and rotate more freely in solution, thus increasing the nuclei T2 relaxation time, leading to the narrower lines^30^ observed in **Figure 1** (top). Permethylation thus allowed the acquisition of NMR data that were useful for structure analysis of the EPS. Importantly, the ^1^H NMR spectrum of **Figure 1** (top), shows three pairs of anomeric signals with a small peak and a large peak with similar intensity ratios of about 1:2. The unequal peak intensities of the pair of signals suggest the possibility of structural irregularity, which is not in agreement with the tetrasaccharide repeating structure predicted from genomic comparison to *X. campestris*.^28^

**Figure 1:**
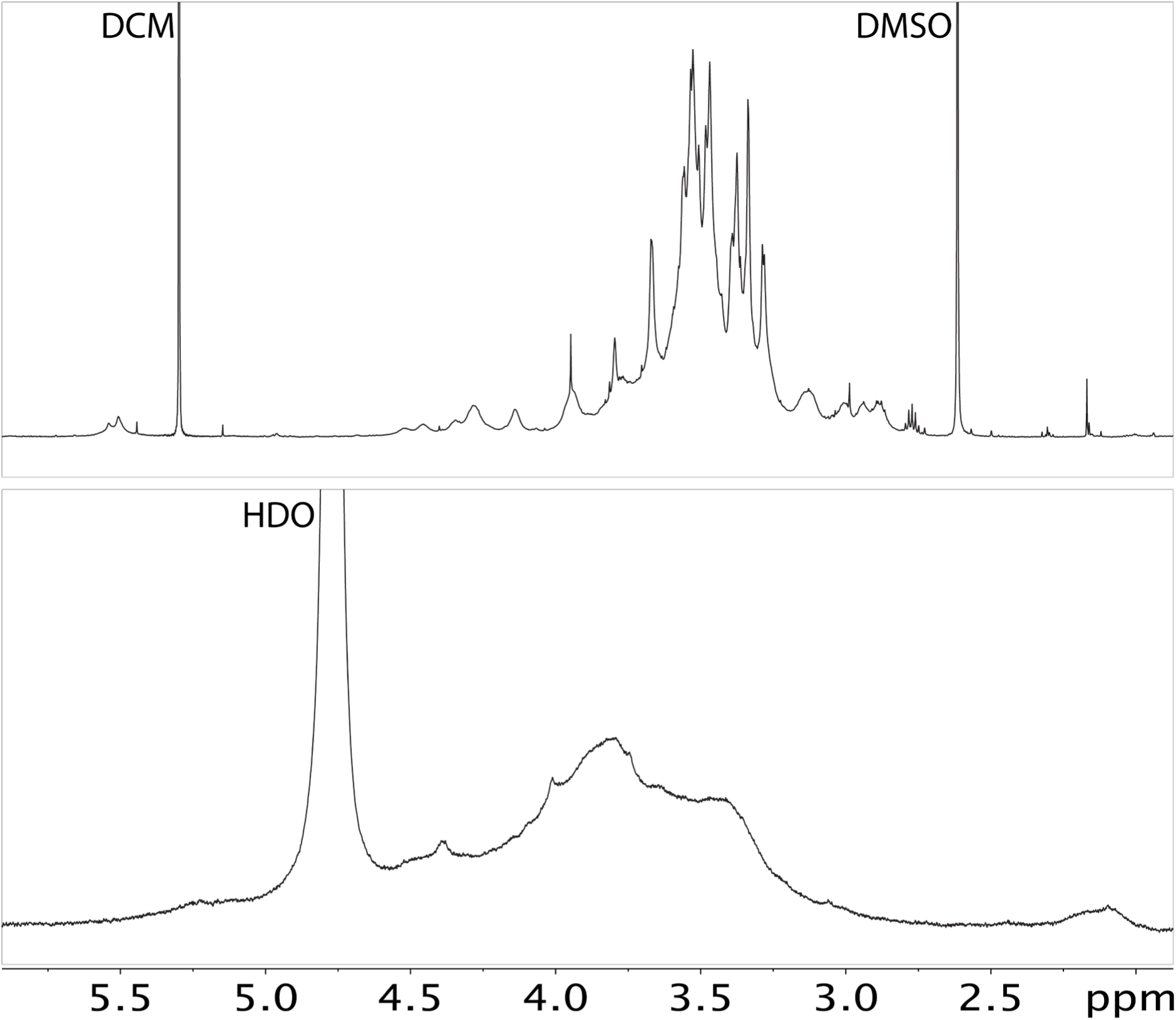
Stacked ^1^H NMR spectra of native *X. fastidiosa* EPS in D_2_O after two rounds of mild acid hydrolysis (bottom) and the permethylated sample acquired in chloroform-*d* (top, solvent peak not shown) – both spectra were acquired at 25 °C. The broad signals of the native sample in D_2_O (bottom) precluded any meaningful analysis. Higher spectral resolution in chloroform-*d* was achieved after permethylation of the sample (top) that revealed salient structure information, including the presence of three pairs of anomeric signals, each of which consisting of a small and a large peak whose intensity ratio was roughly 1:2. The dichloromethane (DCM) and dimethylsulfoxide (DMSO) peaks are from residual solvents in the permethylation reaction.

NMR structure analysis was based on a subset of standard and non-standard 2D experiments, including COSY, HSQC, NOESY, HSQC-TOCSY and HMQC-NOESY that were acquired in lieu of HMBC, which did not yield any observable anomeric correlation. **Table 1** enumerates the spectra utilized for chemical shift assignments for all residues identified for *X. fastidiosa* EPS. For instance, the nuclei connectivity of 3,4-β-Glc*p* residue was established with a combination of NMR experiments, including COSY, TOCSY, HMQC-TOCSY and HMQC-NOESY whereas its anomeric linkage was confirmed with ROESY and HMQC-NOESY data.

**Table 1:**
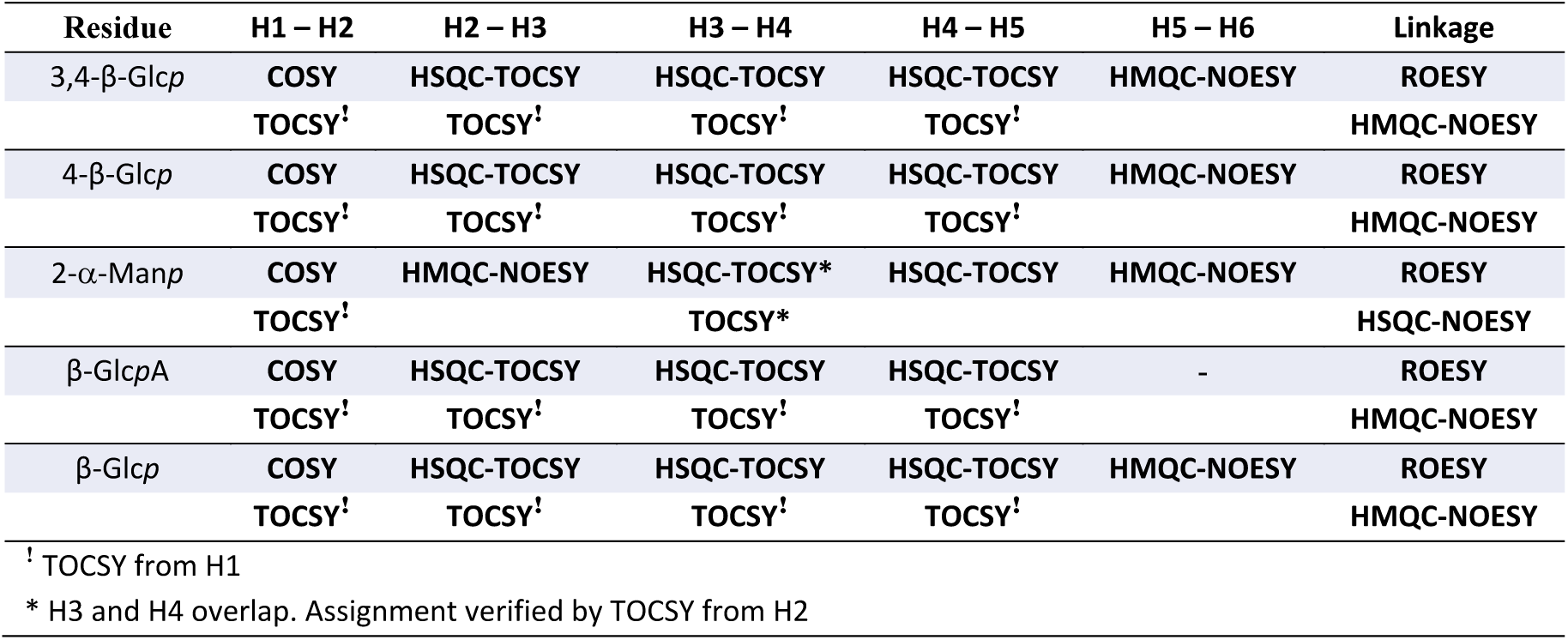
NMR chemical shift assignment protocol for residues identified in *X. fastidiosa* EPS sample.

| Residue | H1 – H2 | H2 – H3 | H3 – H4 | H4 – H5 | H5 – H6 | Linkage |
| --- | --- | --- | --- | --- | --- | --- |
| 3,4-β-Glcp | COSY | HSQC-TOCSY | HSQC-TOCSY | HSQC-TOCSY | HMQC-NOESY | ROESY |
|  | TOCSY<sup>†</sup> | TOCSY<sup>†</sup> | TOCSY<sup>†</sup> | TOCSY<sup>†</sup> |  | HMQC-NOESY |
| 4-β-Glcp | COSY | HSQC-TOCSY | HSQC-TOCSY | HSQC-TOCSY | HMQC-NOESY | ROESY |
|  | TOCSY<sup>†</sup> | TOCSY<sup>†</sup> | TOCSY<sup>†</sup> | TOCSY<sup>†</sup> |  | HMQC-NOESY |
| 2-α-Manp | COSY | HMQC-NOESY | HSQC-TOCSY* | HSQC-TOCSY | HMQC-NOESY | ROESY |
|  | TOCSY<sup>†</sup> |  | TOCSY* |  |  | HSQC-NOESY |
| β-GlcpA | COSY | HSQC-TOCSY | HSQC-TOCSY | HSQC-TOCSY | - | ROESY |
|  | TOCSY<sup>†</sup> | TOCSY<sup>†</sup> | TOCSY<sup>†</sup> | TOCSY<sup>†</sup> |  | HMQC-NOESY |
| β-Glcp | COSY | HSQC-TOCSY | HSQC-TOCSY | HSQC-TOCSY | HMQC-NOESY | ROESY |
|  | TOCSY<sup>†</sup> | TOCSY<sup>†</sup> | TOCSY<sup>†</sup> | TOCSY<sup>†</sup> |  | HMQC-NOESY |

<sup>†</sup> TOCSY from H1
\* H3 and H4 overlap. Assignment verified by TOCSY from H2

NMR spectral analysis, following the protocol in **Table 1**, reveals the presence of two tetrasaccharide sub-structures. Each substructure consists of a 1,4-linked β-Glc*p* disaccharide backbone with a sidechain linked to the 3-position of one β-Glc*p* residue. The sidechain is a disaccharide of α-Man*p* (that is linked to the backbone at the 3-position) substituted at its 2-position by either a terminal β- Glc*p*A or a terminal β-Glc*p* residue. Each monosaccharide residue was identified by its characteristic anomeric ^1^H/^13^C chemical shifts and 2D TOCSY spin-system pattern (**Fig. S4**). In particular, the anomeric protons of Glc*p* and Glc*p*A residues each provide four TOCSY cross peaks that correspond to correlations from H1 to H2, H3, H4 and H5. The latter residue is differentiated from the former by the characteristic deshielding of the H5 signal as well as the methyl ester signal at ∼3.78/51.7 ppm (^1^H/^13^C) – see **Figure 2**. The anomeric protons of the Man*p* residues, on the other hand, provide only one TOCSY cross peak signal from H1 to H2. These residues were also confirmed by methyl alditol glycosyl composition analysis and GC-MS linkage analysis (**Fig. S8 and Tables S2/S3 in Supplemental Information**). The chemical shift values of the assigned residues of *X. fastidiosa* EPS (permethylated) are listed in **Table 2**, while the HSQC signals in **Figure 2** are annotated with the corresponding residue and ring position.

**Table 2:** ^1^H and ^13^C chemical shifts of permethylated *X. fastidiosa* EPS in chloroform-*d* at 45 °C.

| Residue |  | 1 | 2 | 3 | 4 | 5 | 6 |
| --- | --- | --- | --- | --- | --- | --- | --- |
| → 4)-β-Glcp-(1→4 (A') | <sup>1</sup>H | 4.32 | 3.26 | 3.13 | 3.65 | 3.19 | 3.48/3.55 |
|  | <sup>13</sup>C | 102.2 | 83.4 | 85.4 | 79.1 | 75.1 | 72.6 |
| → 4)-β-Glcp-(1→4 (A'') | <sup>1</sup>H | 4.30 | 3.25 | 3.12 | 3.69 | 3.16 | 3.47/3.57 |
|  | <sup>13</sup>C | 102.4 | 83.7 | 85.4 | 79.4 | 74.9 | 72.6 |
| → 3,4)-β-Glcp-(1→4 (B) | <sup>1</sup>H | 4.38 | 2.90 | 3.75 | 3.98 | 3.45 | 3.68/3.73 |
|  | <sup>13</sup>C | 104.2 | 82.6 | 76.0 | 74.9 | 74.0 | 70.2 |
| → 3,4)-β-Glcp-(1→4 (C) | <sup>1</sup>H | 4.30 | 2.89 | 3.74 | 3.96 | 3.30 | 3.71/3.71 |
|  | <sup>13</sup>C | 104.0 | 82.6 | 76.1 | 74.9 | 74.2 | 69.9 |
| → 2)-α-Manp-(1→3 (D) | <sup>1</sup>H | 5.53 | 4.15 | 3.48 | 3.47 | 3.97 | 3.50/3.60 |
|  | <sup>13</sup>C | 98.7 | 76.4 | 80.0 | 76.3 | 70.9 | 72.4 |
| → 2)-α-Manp-(1→3 (E) | <sup>1</sup>H | 5.50 | 4.15 | 3.45 | 3.47 | 3.95 | 3.50/3.60 |
|  | <sup>13</sup>C | 98.8 | 76.2 | 80.0 | 76.3 | 70.7 | 72.4 |
| β-GlcpA-(1→4 (F) | <sup>1</sup>H | 4.52 | 3.00 | 3.31 | 3.36 | 3.82 | - |
|  | <sup>13</sup>C | 104.3 | 83.7 | 85.9 | 81.1 | 73.8 | 169.4 |
| β-Glcp-(1→4 (G) | <sup>1</sup>H | 4.43 | 2.94 | 3.27 | 3.01 | 3.35 | 3.58/3.58 |
|  | <sup>13</sup>C | 104.0 | 84.1 | 86.4 | 80.3 | 74.6 | 72.9 |

**Figure 2:**
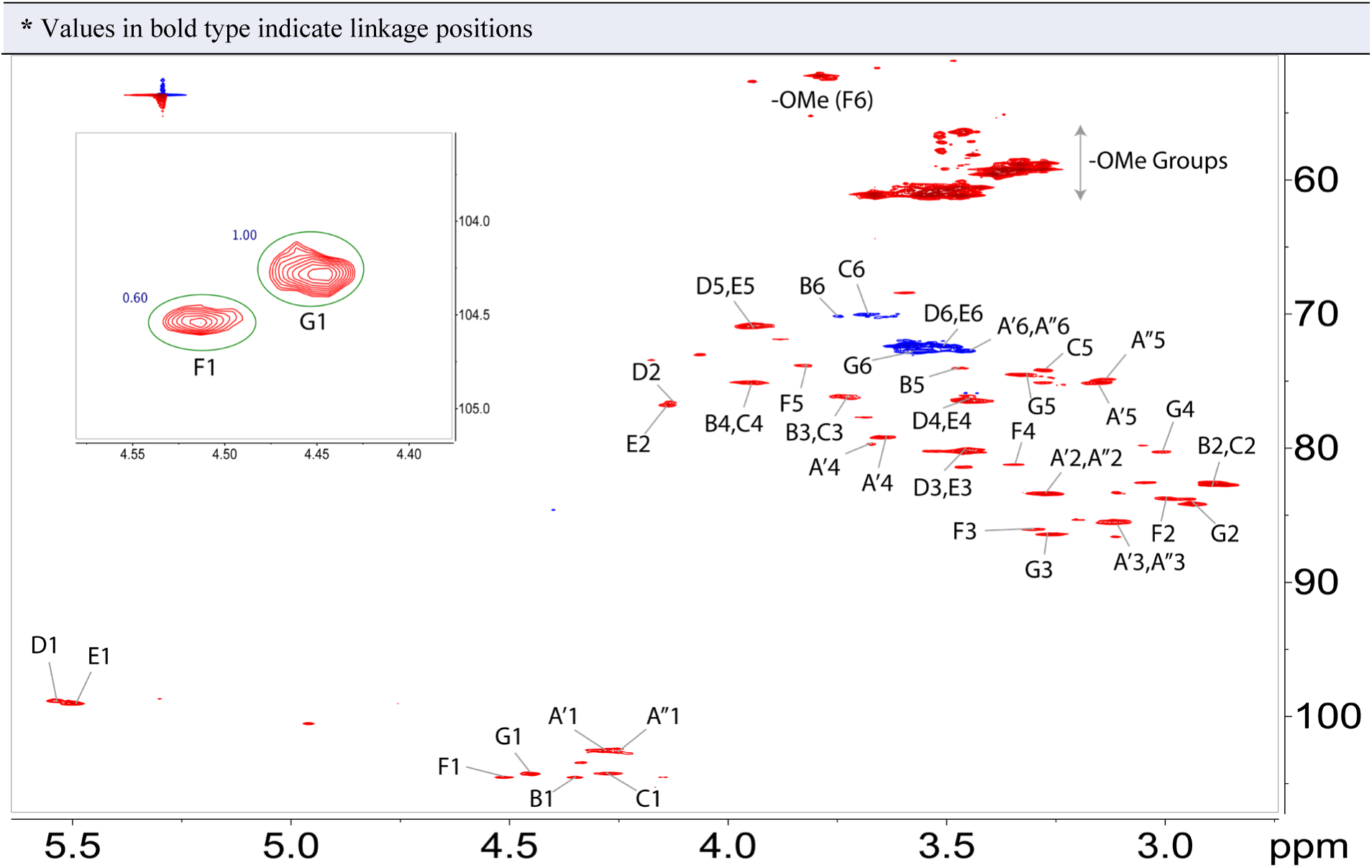
Partial ^1^H-^13^C-HSQC NMR spectrum of permethylated *X. fastidiosa* EPS, acquired at 45 °C. Residue signals of the EPS are annotated in addition to monosaccharide signals (circled and annotated M) generated after mild acid hydrolysis of the permethylated sample. Signals not annotated are either from the monosaccharides generated or unassigned/unrelated residue. Inset is the expansion of H1 signals of residues G and F, showing the relative peak intensity ratio of G : F (1: 0.6).

The pairs of anomeric signals shown in **Figure 1** (top) correspond to signals from the two tetrasaccharide units shown below;

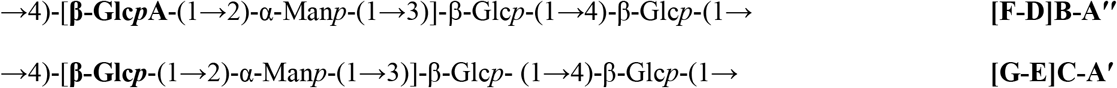

The pair of tetrasaccharides are not equimolar, as is evident from the unequal proton peak intensities in **Figure 1** (top). The pair of tetrasaccharide units were linked through Residues **A′** and **A′′**, and the observation of weak long-range inter-residue ROESY signals from **D1** → **A′2, E1** → **A′2** and **E1** → **A′′2**, were vital to elaborate the sequence of these two building blocks (**Figure 3** left). These ROESY signals confirmed the following anomeric linkages; **A′1** → **B4, A′1** → **C4** and **A′′1** → **C4**, respectively. In particular, the absence of ROESY signal from **D1** → **A′′2** indicated the absence of a linkage from **A′′1** → **B4** in the polysaccharide backbone structure. With the foregoing ROESY information at hand, the polysaccharide backbone linkage could now be deduced from the overlapping HMQC-NOESY correlation peak shape, shown in **Figure 3** (right), thus leading to the proposed EPS structure of *X. fastidiosa*, shown in **Figure 4**. The repetition rate ‘n’ and ‘m’ can be deduced from the HSQC anomeric peak integrals of residues **G** : **F** (inset **Figure 2**), shown to be ∼1 : 0.6, which also corresponds to the ratios of the two tetrasaccharides. The ratio n : m is thus calculated to be 3 : 2.

**Figure 3:**
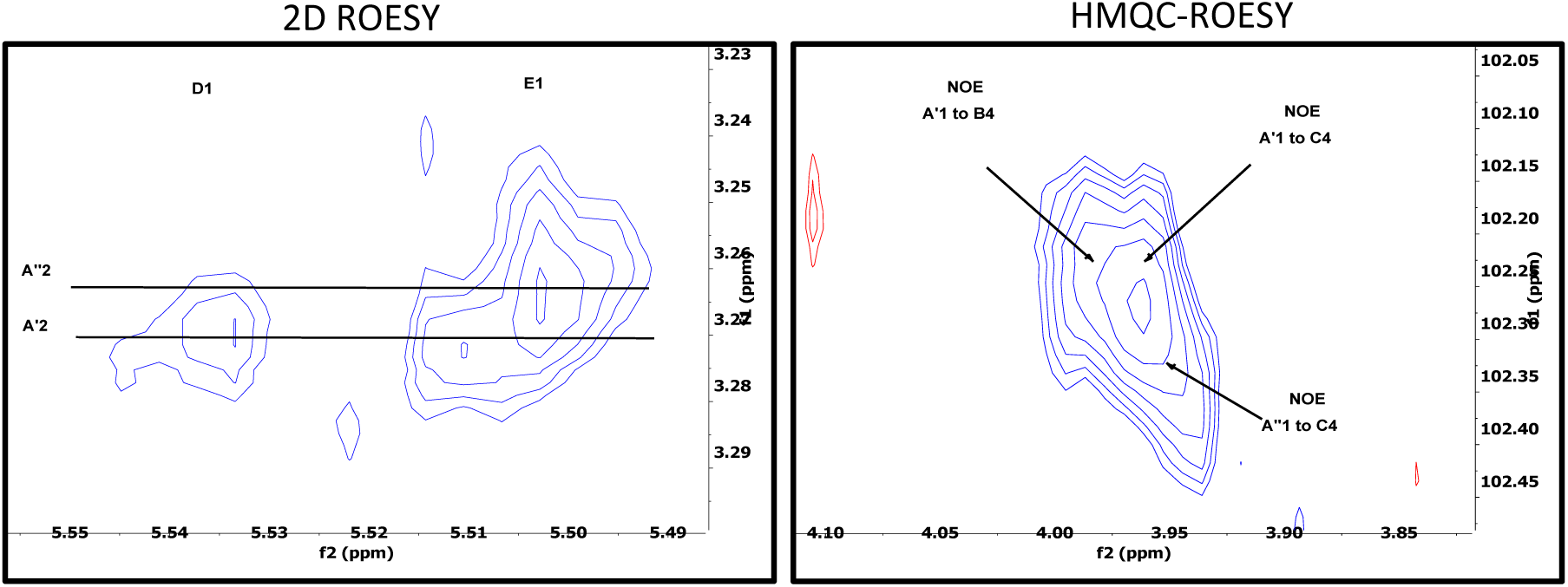
Expansion of important ROESY and HMQC-ROESY correlation peaks, showing inter-residue connectivity between D1 and A’2 as well as from E1 to both A’2 and A’’2 (left spectrum). The absence of ROESY correlation from D1 to A’’2 was useful for disambiguation of the HMQC-ROESY correlation peaks (right spectrum) that shows the order of backbone linkage of *X. fastiosa* EPS. Note that the number notation refers to the proton nuclei of interest (that is, D1 refers to proton-1 of residue D).

**Figure 4:**
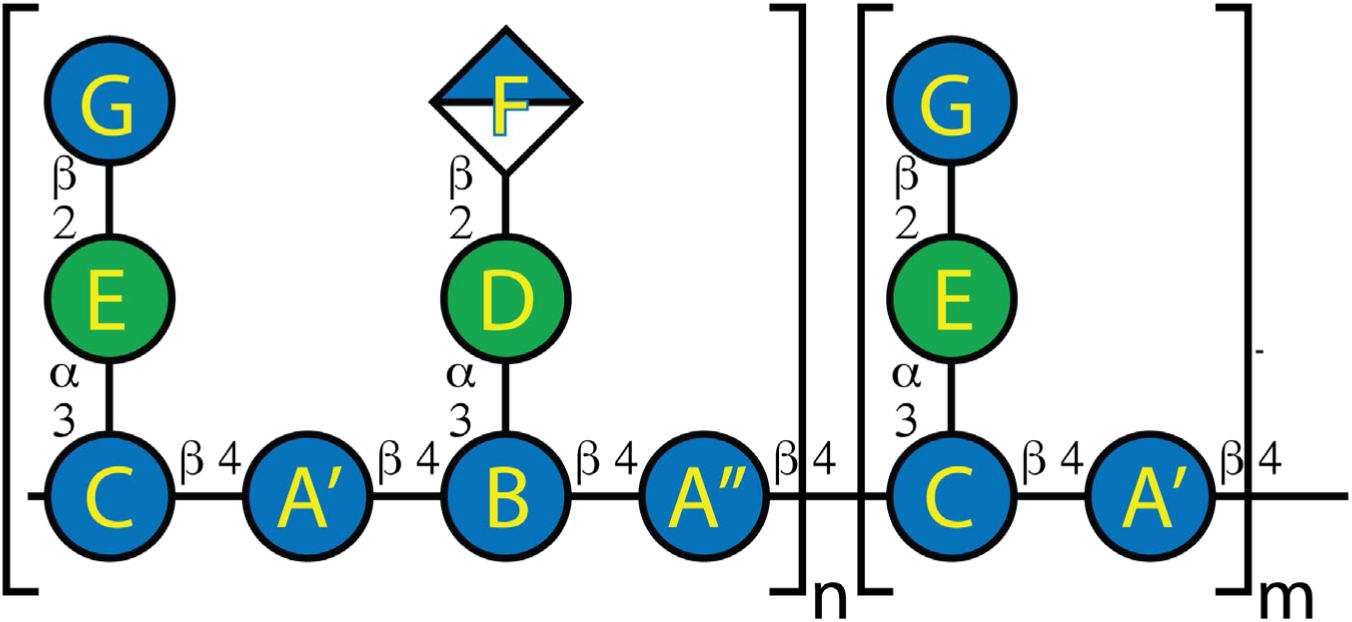
The repeating octasaccharide structure of the native *X. fastidiosa* EPS – note that the permethylated sample will have -OMe groups in place of -OH groups. The residues are labelled A′, A′′, B, C, D, E, F, and G to match the residues in the chemical shift table (**Table 2**). Glc*p*, Man*p* and Glc*p*A residues are represented with blue circle, green circle and blue/white diamond shapes. The ratio of n : m is determined as 3 : 2 based on HSQC anomeric signals integration (**Figure 2**).

## Conclusion

The limitations inherent in the utilization of solution-state NMR spectroscopy for structure characterization of high molecular weight polysaccharides, can be ameliorated by permethylation strategies. Polysaccharide permethylation allows their solubilization in less viscous solvents like chloroform-*d* but does not lead to significant changes in the well-established chemical shift region definition of residue nuclei or the salient distinction between different monosaccharides. In particular, 2D TOCSY NMR spin-lock pattern and the ring residue unique anomeric NMR chemical shifts can be used to differentiate these monosaccharides – making the permethylation approach useful for structure characterization protocols. Application of the permethylation strategy, in combination with other analogous methods, allowed an effective structure analysis of the exopolysaccharide of *X. fastidiosa*. The *X. fastidiosa* EPS was determined to consist of a cellobiose backbone with 3-linked sidechains on alternating glucose units of 1,2-α-mannose terminating with either β-glucuronic acid or β-glucose residue.

## Acknowledgement

This work was supported by the U.S. Department of Energy, Office of Science, Basic Energy Sciences, grant number DE-SC0015662 and by the NIH General Medical Sciences grant number R24GM137782 to Parastoo Azadi.

