## Supplemental Information for "Permethylation as a Strategy for High Molecular Weight Polysaccharide Structure Analysis by NMR – Case Study of *Xylella fastidiosa* EPS"

#### Experimental

##### NMR Acquisition:

NMR data were acquired on a Bruker 600 MHz spectrometer equipped with a 5 mm TXI cryo-probe at probe temperature 45 ºC, except otherwise stated. Between 5 - 10 mg of the samples were permethylated and dissolved in 550 µL chloroform-*d* prior to NMR acquisition. The following general parameters were used for NMR data acquisition. Specific parameters used are indicated in the captions of the respective spectra.

***Xylella fastidiosa***: $^1$H NMR experiment (600 Hz spectral width, 8,192 t1 points and 4 scans); COSY experiment (5144.0*5144.0 F1*F2 spectral width, 1024*512 t2*t1 points and 4 scans); HSQC experiment (5144.0*10570.8 F1*F2 spectral width, 1024*512 t2*t1 points, 145 Hz $^1J_{CH}$ delay and 10 scans); HSQC_TOCSY experiment (5208.3*10570.8 F1*F2 spectral width, 1024*512 t2*t1 points, 145 Hz $^1J_{CH}$ delay, 18 ms mixing time and 24 scans); HMQC_NOESY experiment (5208.3*10570.8 F1*F2 spectral width, 1024*512 t2*t1 points, 145 Hz $^1J_{CH}$ delay, 60 ms mixing time and 182 scans); HMBC experiment (5319.1*10570.8 F1*F2 spectral width, 1536*512 t2*t1 points at 50 % non-uniform sampling in the indirect dimension, 8 Hz $^1J_{CH}$ delay and 540 scans); TOCSY experiment (6250.0*6250.0 F1*F2 spectral width, 1536*512 t2*t1 points, 60 ms spin-lock time and 8 scans); ROESY experiment (6250.0*6250.0 F1*F2 spectral width, 1536*512 t2*t1 points, 70 ms mixing time and 24 scans).

***TFA hydrolyzate of intact EPS***: Quantitative $^1$H NMR spectrum of TFA-hydrolyzed EPS was acquired at 25 °C using 11900 Hz spectral width, 16384 points, four scans and a total relaxation delay of 63 s between each scan. The reference spectra of glucurono-6,3-lactone and glucose/glucuronic acid mixed standard were acquired with the same parameters.

### *X. fastidiosa* EPS assigned impurity

**Table 1:** $^{1}$H and $^{13}$C chemical shifts of rhamnose impurity in permethylated *X. fastidiosa* EPS sample acquired in chloroform-*d* at 45 ºC.

| Residue | | 1 | 2 | 3 | 4 | 5 | 6 |
|---|---|---|---|---|---|---|---|
| → 2)-α-Rha*p*-(1→2 **(H)** | $^{1}$H | **4.96** | **4.06** | 3.46 | 3.04 | 3.60 | 1.25 |
| | $^{13}$C | **100.5** | **73.0** | 81.4 | 82.6 | 68.4 | 18.0 |
| * Values in bold type indicate linkage positions | | | | | | | |

#### *X. Fastidiosa* EPS NMR Data

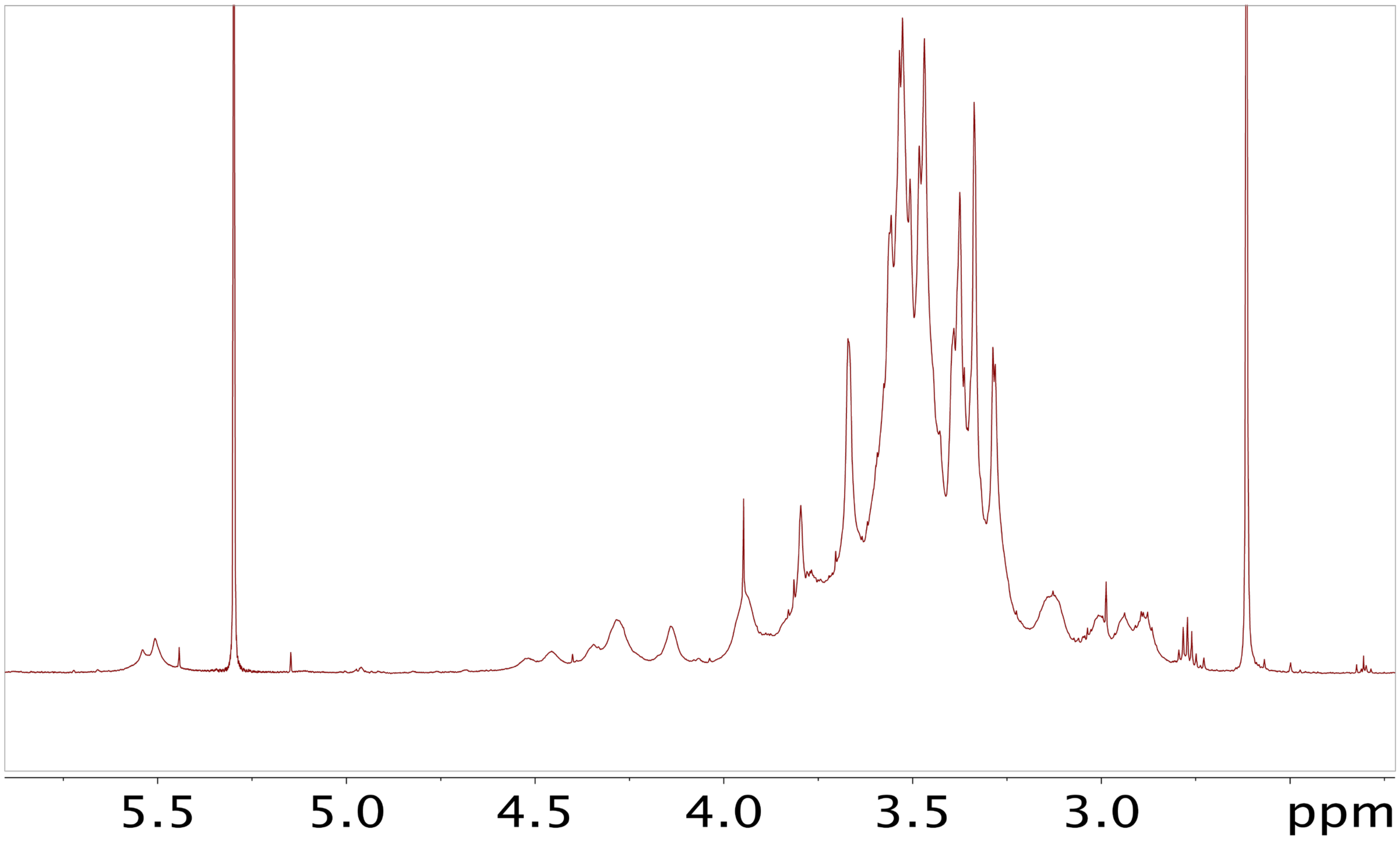


**Figure S1:** $^{1}$H NMR spectrum of permethylated *X. fastidiosa* EPS acquired on a Bruker 600 MHz spectrometer equipped with 5 mm TXI cryo-probe at 25 ºC.

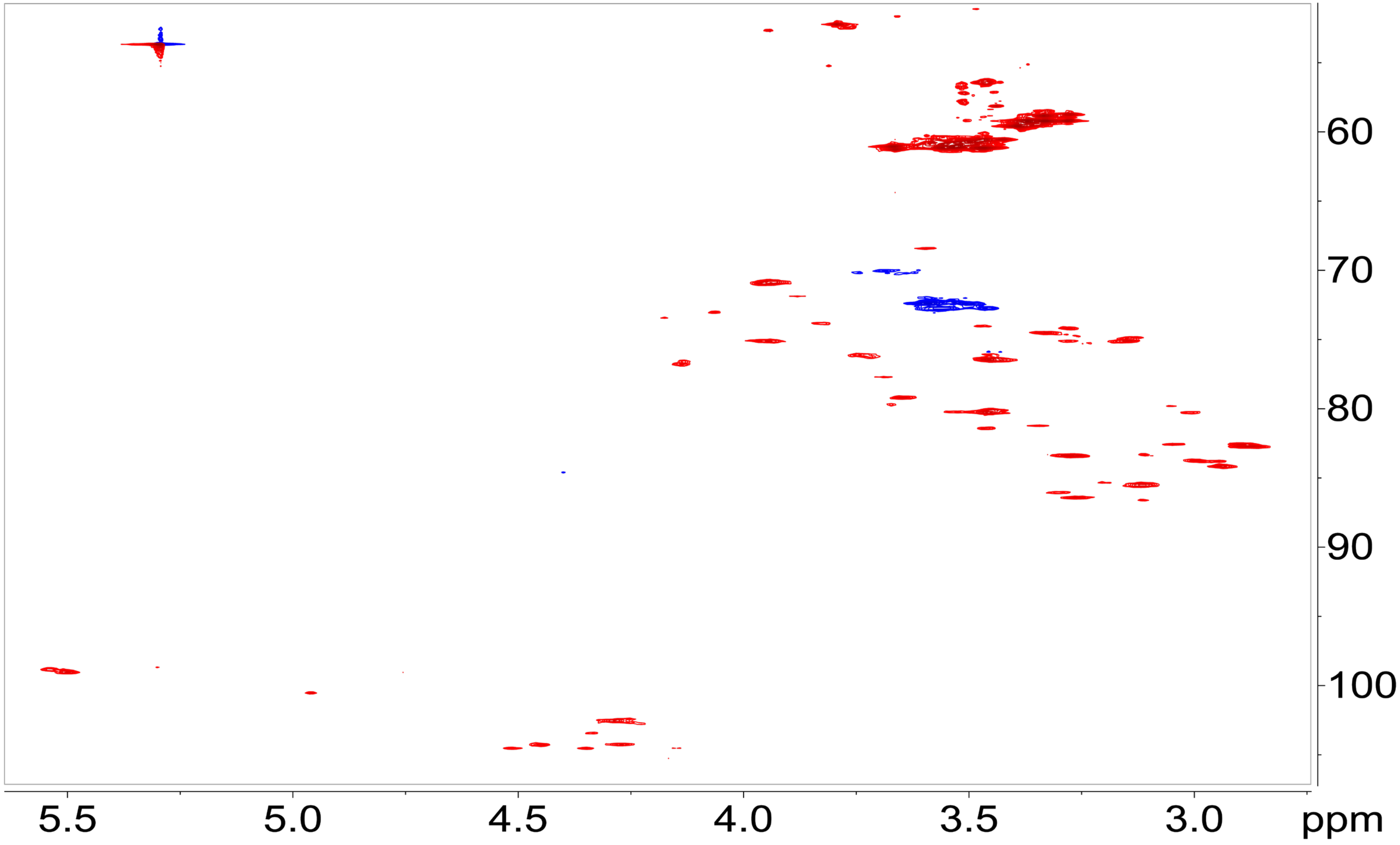


**Figure S2:** $^{1}H$-$^{13}C$ HSQC NMR spectrum of permethylated *X. fastidiosa* EPS acquired on a Bruker 600 MHz spectrometer equipped with 5 mm TXI cryo-probe at 25 ºC, with 145 Hz $^{1}J_{CH}$ delay.

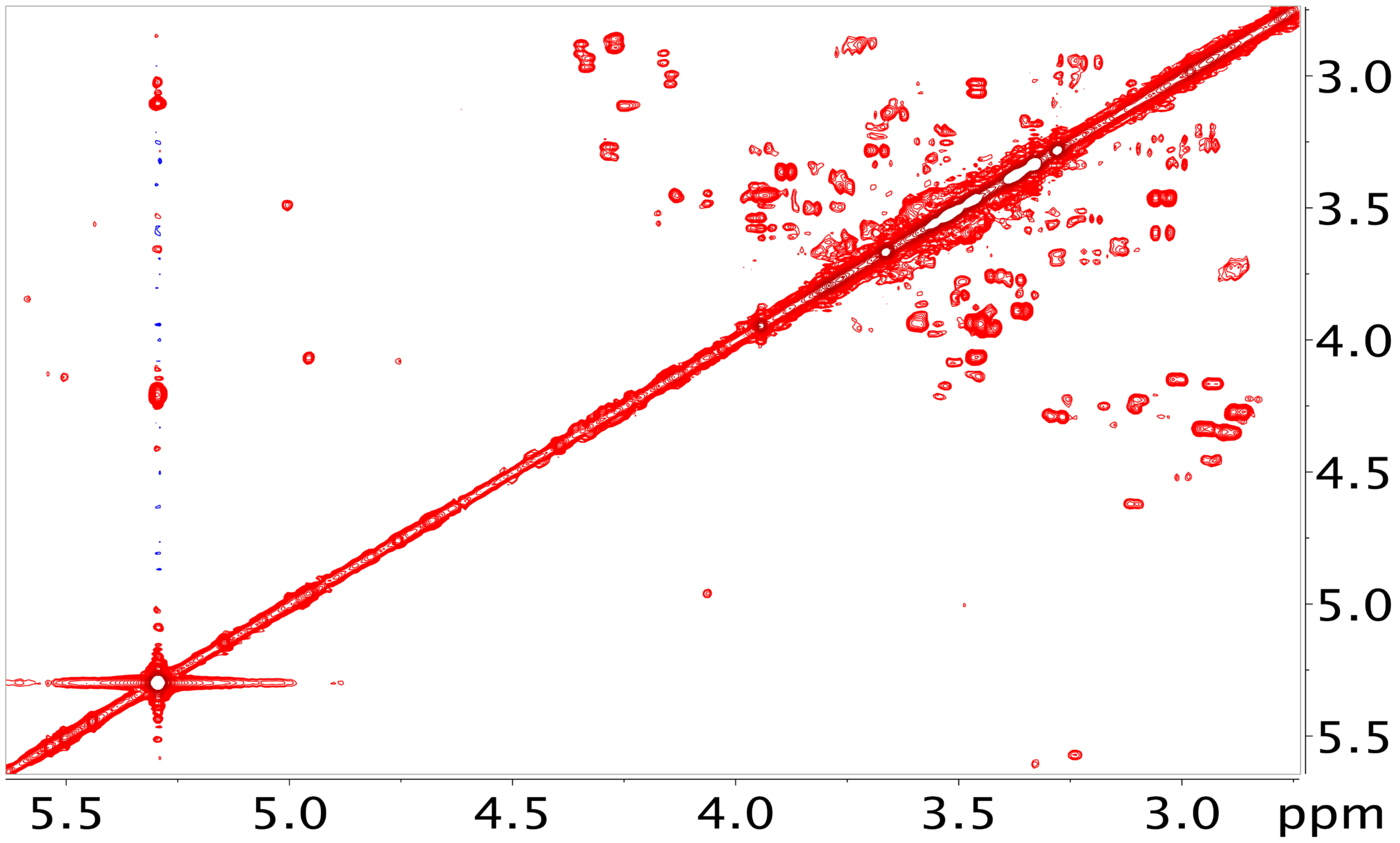


**Figure S3:** $^{1}$H-$^{1}$H COSY NMR spectrum of permethylated *X. fastidiosa* EPS acquired on a Bruker 600 MHz spectrometer equipped with 5 mm TXI cryo-probe at 25 ºC.

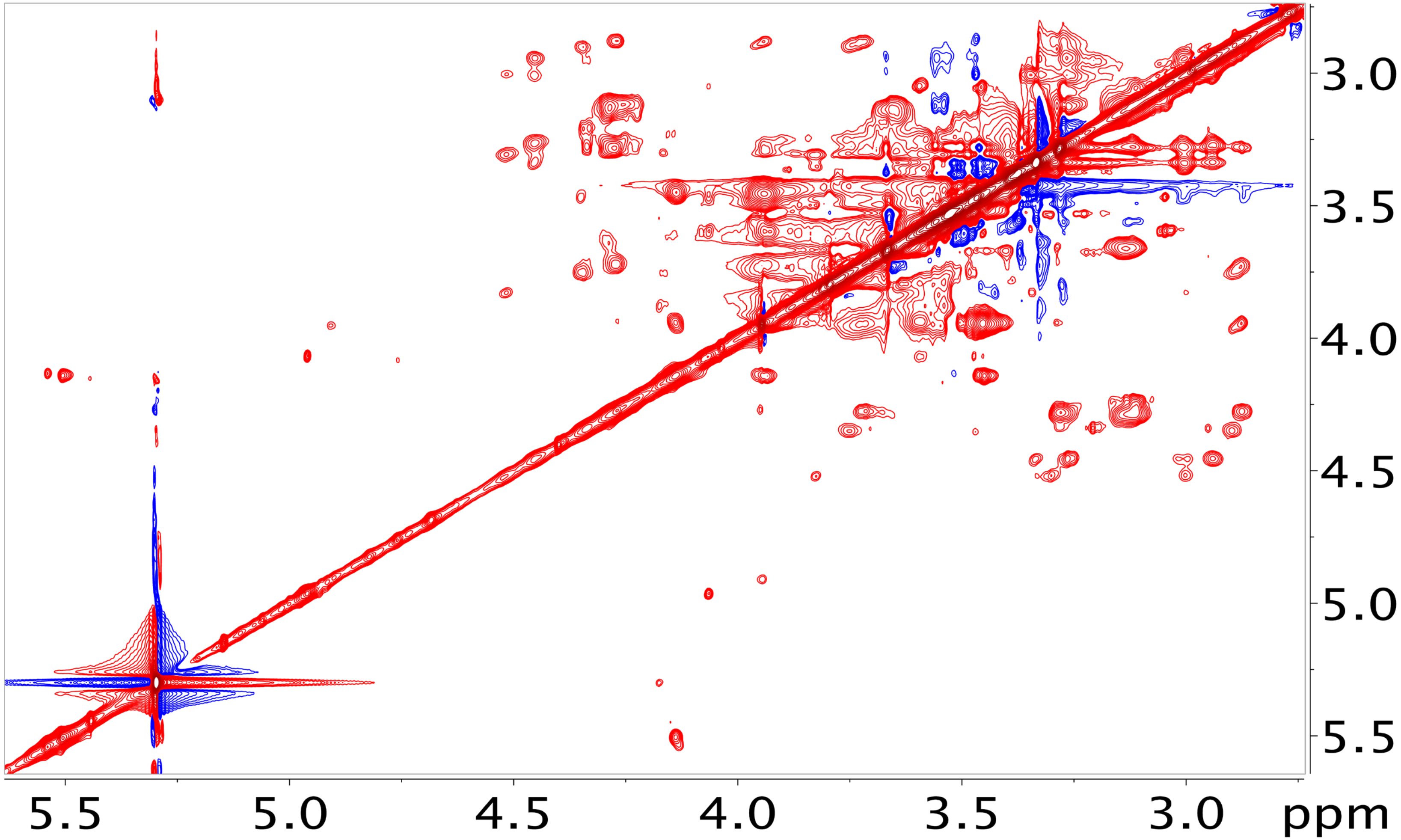


**Figure S4:** $^{1}H$-$^{1}H$ TOCSY NMR spectrum of permethylated *X. fastidiosa* EPS acquired on a Bruker 600 MHz spectrometer equipped with 5 mm TXI cryo-probe at 45 ºC, with 70 ms mixing time.

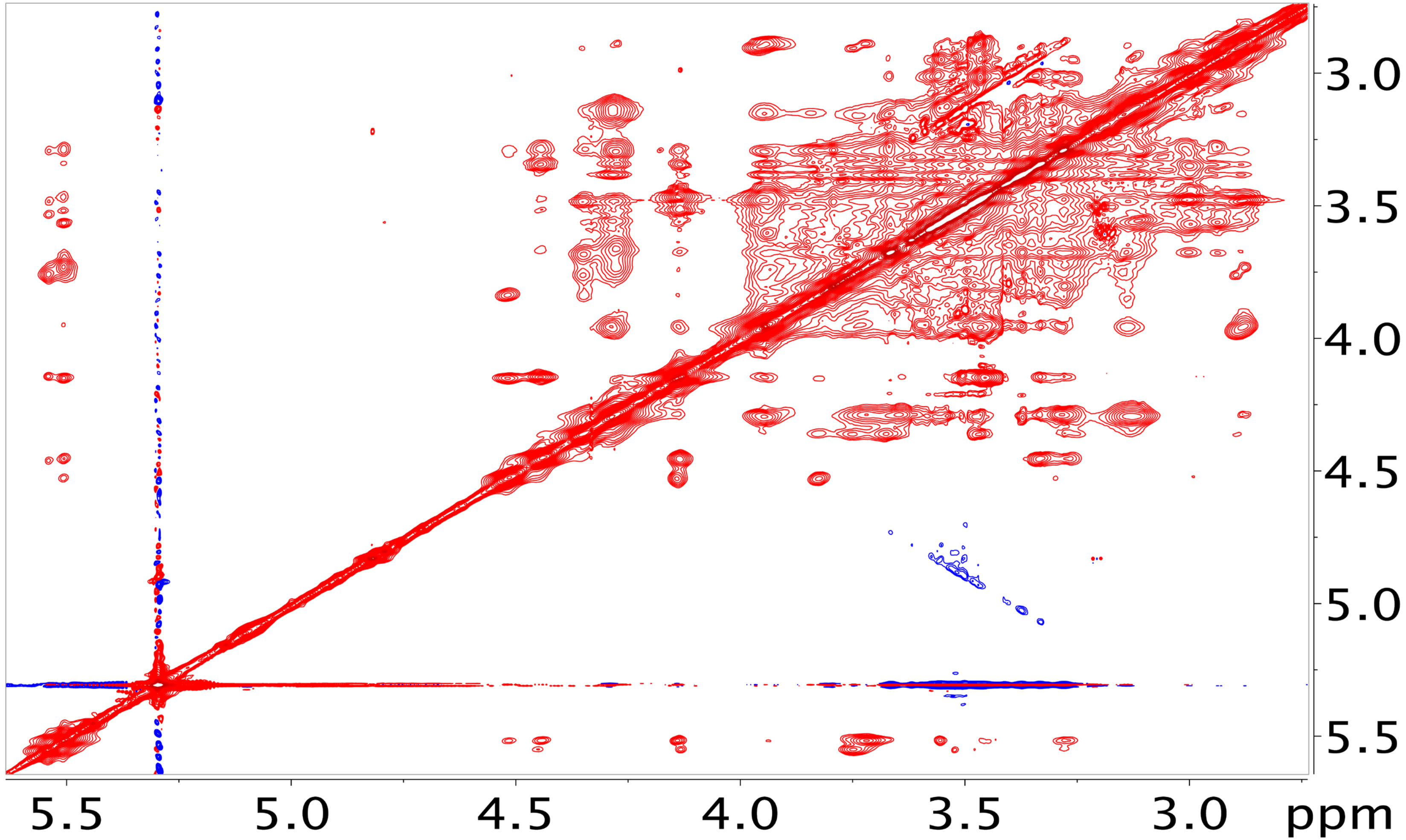


**Figure S5:** $^{1}H$-$^{1}H$ NOESY NMR spectrum of permethylated *X. fastidiosa* EPS, acquired on a Bruker 600 MHz spectrometer equipped with 5 mm TXI cryo-probe at 45 ºC.

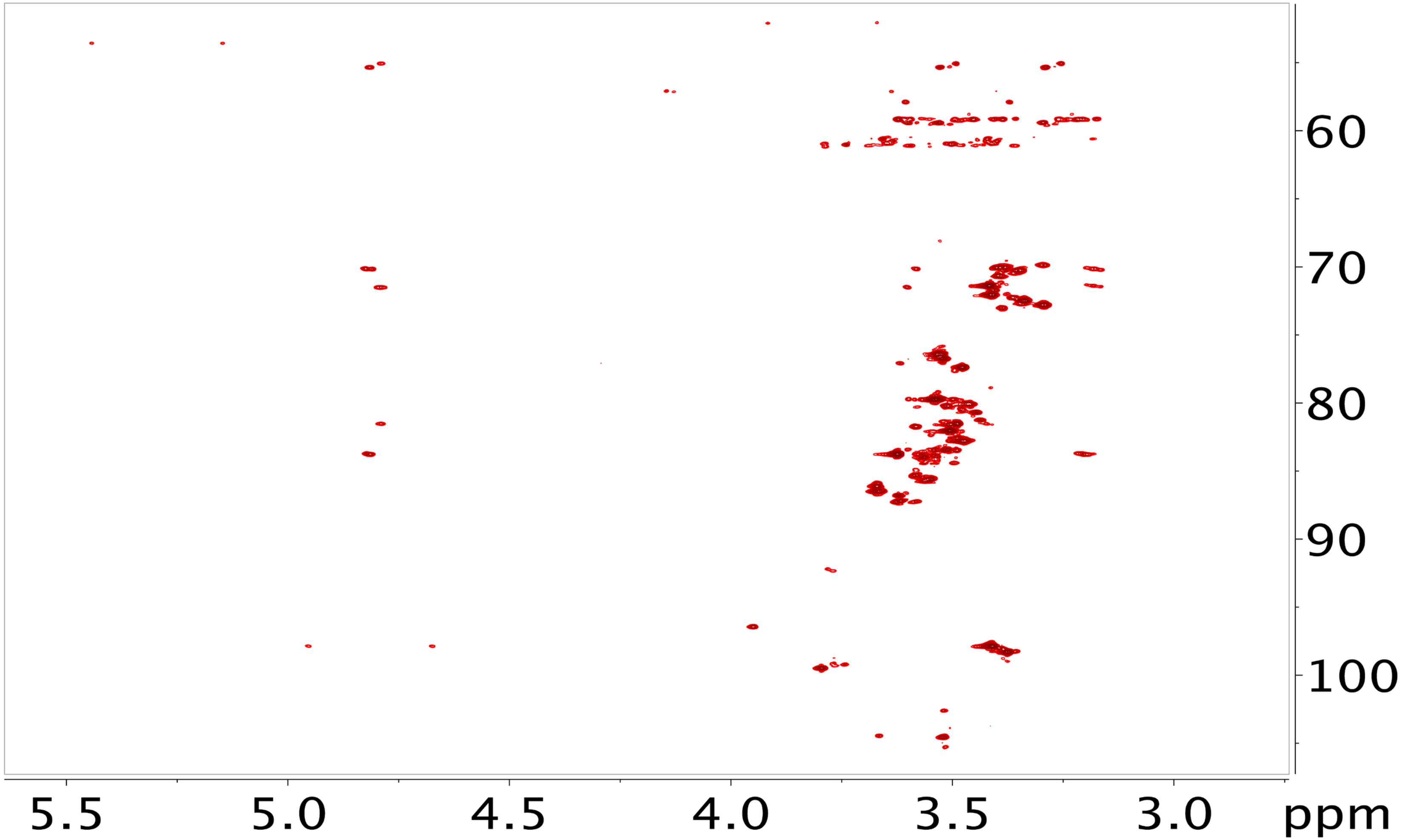


**Figure S6:** $^{1}H$-$^{13}C$ HMBC NMR spectrum of permethylated *X. fastidiosa* EPS, acquired on a Bruker 600 MHz spectrometer equipped with 5 mm TXI cryo-probe at 45 ºC.

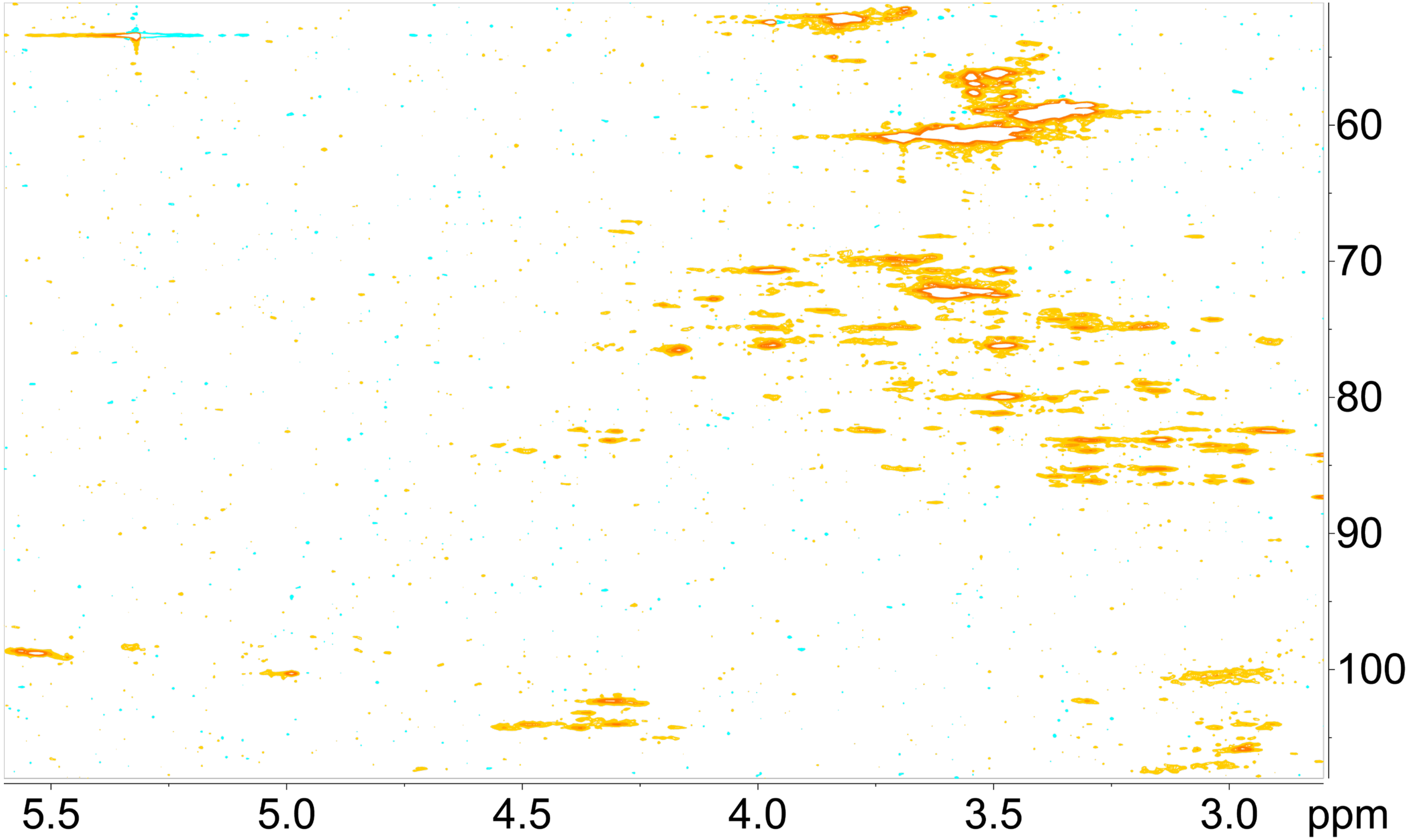


**Figure S7**: $^{1}H$-$^{13}C$ HSQC-TOCSY NMR spectrum of permethylated *X. fastidiosa* EPS, acquired on a Bruker 600 MHz spectrometer equipped with 5 mm TXI cryo-probe at 45 ºC.

### Glycosyl composition analysis by methyl alditol (MA) method

Glycosyl composition analysis was performed by combined gas chromatography/mass spectrometry (GC/MS) of the per-*O*-methyl alditol (MA) derivatives of the monosaccharides produced from the sample as described previously by Black *et al.* (2019) *Analytical Chemistry* **91, 21**: 13787-13793, with slight modification.

Composition analysis of an aliquot of the NMR methylated sample was performed. The methylated sample was hydrolyzed using 2 M TFA (2 h in sealed tube at 121 °C) and reduced with $NaBD_4$. After drying, the sample was re-permethylated to derivatize the hydroxyls created during the hydrolysis. The resulting MAs were analyzed on an Agilent 7890A GC interfaced to a 5975C MSD (mass selective detector, electron impact ionization mode); separation was performed on a 30 m Supelco Equity-1 bonded phase fused silica capillary column.

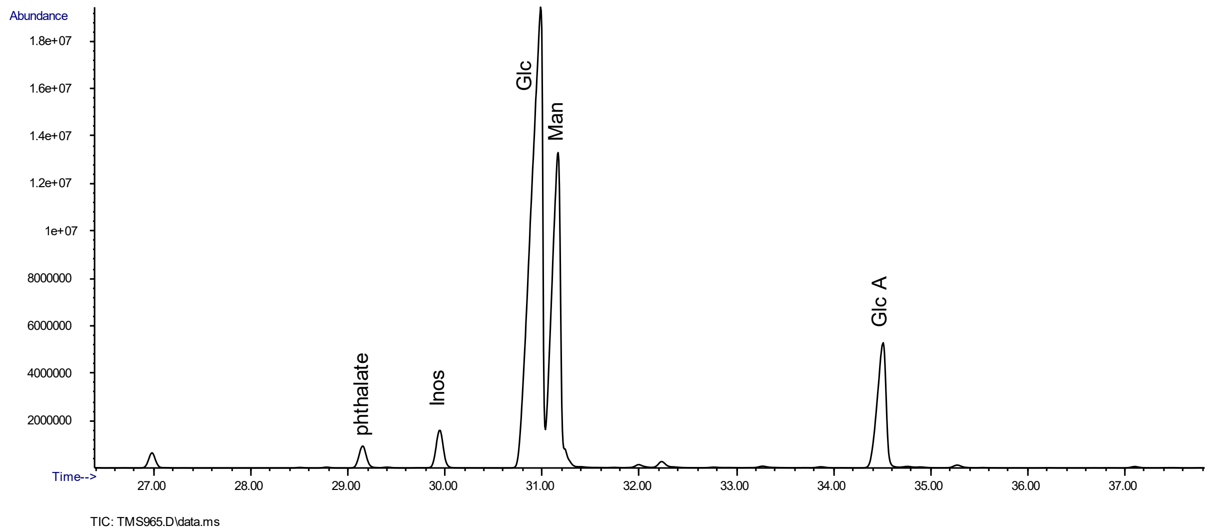


**Figure S8:** Glycosyl composition analysis of *X. fastidiosa* EPS by the MA method.

### Glycosyl linkage analysis by methylation

Glycosyl linkage analysis was performed by combined gas chromatography/mass spectrometry (GC/MS) of the partially methylated alditol acetates (PMAAs) derivatives produced form the samples. The procedure is a slight modification of the one described by Willis *et al.* (2013) *PNAS*, 110 (19) 7868-7873.

Linkage analysis of an aliquot of the NMR methylated sample was performed. The methyl esterified glucuronic acids were reduced using $LiAlD_4$ (300 µl, 10 mg/ml) in THF for 8 hours at 80 °C. The reduced sample was then hydrolyzed using 2 M TFA (2 h in sealed tube at 120 °C), reduced with $NaBD_4$, and acetylated using acetic anhydride/TFA. The resulting PMAAs were analyzed on an Agilent 7890A GC interfaced to a 5975C MSD (mass selective detector, electron impact ionization mode); separation was performed on a 30 m Supelco SP-2331 bonded phase fused silica capillary column.

**Table S2:** Amount and mole percentage of each residue detected in the *X. fastidiosa* EPS by the MA method.

| Glycosyl residue | Area % |
|---|---|
| Mannose (Man) | 29.8 |
| Glucose (Glc) | 58.3 |
| Glucuronic Acid (Glc A) | 11.9 |

**Table S3:** Relative ratios of the partially methylated alditol acetate (PMAA) peaks observed in the linkage analysis.

| Residue | Area % |
|---|---|
| Terminal mannopyranosyl residue (t-Man) | 1.6 |
| Terminal glucopyranosyl residue (t-Glc) | 14.0 |
| 2 linked mannopyranosyl residue (2-Man) | 24.0 |
| Terminal glucopyranosyl uronic acid residue (t-Glc A) | 7.0 |
| 4 linked glucopyranosyl residue (4-Glc) | 29.8 |
| 3,4 linked glucopyranosyl residue (3,4-Glc) | 23.6 |

#### Glycosyl composition by quantitative $^{1}$H NMR

Intact EPS (~5 mg) was completely hydrolyzed by treatment with 2 M TFA (2 h in sealed tube at 120 °C) and the TFA was evaporated with a stream of nitrogen. The dry sample was deuterium-exchanged by lyophilizing from $D_2O$ (99.9% D, Sigma) twice. The dry material was dissolved in 510 µL $D_2O$ (99.96% D, Cambridge Isotope), 50 nmol DSS-$d_6$ chemical shift standard were added and the sample was transferred into a 5 mm NMR tube. The pH of the sample (nominal reading) was measured using a combined micro-electrode (Accumet). A sample of glucurono-6,3-lactone was prepared by dissolving 2 mg of the material (Sigma) in 510 µL $D_2O$ (99.9% D) with added 50 nmol DSS-$d_6$. A mixed sample of glucose and glucuronic acid was prepared by dissolving ~3.5 mg of each monosaccharide (Sigma) in 510 µL $D_2O$ (99.9% D) with added 50 nmol DSS-$d_6$. The pH of this sample was adjusted with sodium deuteroxide (Sigma) so that the pH reading was the same as for the TFA-hydrolyzed EPS sample (pH 3.2).

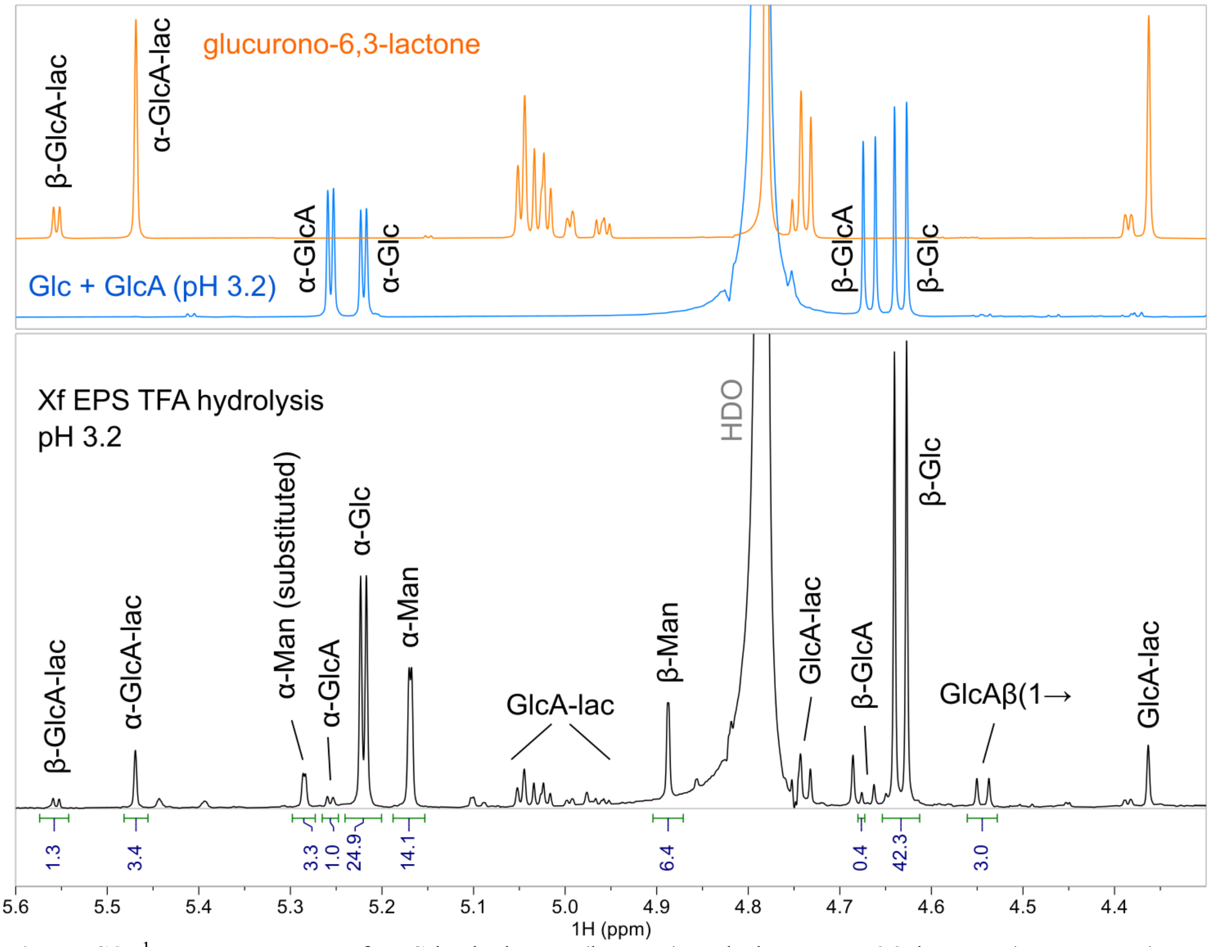


**Figure S8:** $^{1}$H NMR spectra of EPS hydrolyzate (bottom) and glucurono-6,3-lactone (top, orange) and a mixed standard of glucose and glucuronic acid (top, blue). Assignments of signals in the anomeric region are indicated.

$^{1}$H NMR spectrum of the EPS hydrolyzate (Fig. S8) contained signals in the anomeric region that belonged to α/β-Glc, α/β-Man, and α/β-GlcA. Also present were signals of α/β-glucurono-6,3-lactone that formed from GlcA during the hydrolysis. Two additional signals of approximately the same intensities were present: (i) The doublet (8.0 Hz) at 4.54 ppm was assigned as a terminal β-GlcA, based on sensitivity of its position on pH changes (not shown), and (ii) the doublet (1.3 Hz) at 5.29 ppm was assigned as a substituted α-Man. Combined with the structural findings on the permethylated EPS, it is likely that these two residues form a residual GlcA*β*(1→2)Man*α* disaccharide that was resistant to the hydrolysis. Additional minor signals present in the spectrum were not identified.

The populations of Glc, Man and GlcA were determined based on combined integrals of signals that represent each monosaccharide: Glc (α+β Glc), Man (α+β Man, α-Man substituted), GlcA (α+β GlcA, α+β GlcA-lac, GlcAβ linked). The experimentally determined ratio of Glc:Man:GlcA is thus 67:24:9, whereas the theoretical ratio is 66:25:9 for the proposed EPS structure where n:m = 3:2.
